# Sex-linked gene expression and the reversion to hermaphroditism in *Carica papaya* L. (Caricaceae)

**DOI:** 10.1101/2020.06.25.169623

**Authors:** T Chae, A Harkess, RC Moore

## Abstract

One evolutionary path from hermaphroditism to dioecy is via a gynodioecious intermediate. The evolution of dioecy may also coincide with the formation of sex chromosomes that possess sex-determining loci that are physically linked in a region of suppressed recombination. Dioecious papaya (*Carica papaya*) has an XY chromosome system, where the presence of a Y chromosome determines males. However, in cultivation, papaya is gynodioecious, due to the conversion of the male Y chromosome to a hermaphroditic Y^h^ chromosome during its domestication. We investigated gene expression linked to the X, Y, and Y^h^ chromosomes at different floral developmental stages in order to identify differentially expressed genes (DEGs) that may be involved in the sexual reversion of males to hermaphrodites. We identified 309 sex-biased genes found on the sex chromosomes, most of which are found in the pseudoautosomal regions (PARs). Female (XX) expression in the sex determining region (SDR) was almost double that of X-linked expression in males (XY) and hermaphrodites (XY^h^), which rules out dosage compensation for most sex-linked gene; although, an analysis of hemizygous X-linked loci found evidence of partial dosage compensation. Furthermore, we identified a potential candidate gene associated with both sex determination and the transition to hermaphroditism, a homolog of the MADS-box protein *SHORT VEGETATIVE PHASE* (*SVG*).

## INTRODUCTION

Over 90% of flowering plant species are hermaphroditic, which is beneficial in circumstances where selfing is necessary for reproductive assurance (Renner, 2014). In contrast, only 6% of flowering plant species are dioecious, with individually staminate (male) or pistillate (female) flowers (Renner, 2014). Although dioecy is uncommon, it is predicted to be evolutionarily advantageous as it promotes outcrossing, which increases heterozygosity and adaptability (Barrett, 2013; Glick et al., 2016). Despite its low frequency, dioecy is represented in approximately half of all flowering plant families (Barrett, 2013; Charlesworth, 2018; Henry et al., 2018) and has evolved independently multiple times. Transitioning from hermaphroditism to dioecy includes the benefits of inbreeding avoidance and resource reallocation. This transition leads to changes that may maximize developmental and physiological traits more beneficial in separate sexes than in a cosexual state (Charlesworth, 2018). This ‘trade-off’ hypothesis would lead to major adaptations of females and males and enhance their female and male functions, respectively (Charlesworth, 2018).

One path to the evolution of dioecy involves the emergence of a genetic regulatory mechanism using sex chromosomes (Moore et al., 2016). One model that explains the emergence of sex chromosomes involves mutations at two fertility loci (Harkess et al., 2017; Charlesworth, 2018; Akagi et al., 2019; Harkess et al., 2020). The first mutation typically is recessive and causes male sterility, leading to gynodioecious populations of pistillate individuals and hermaphrodites (Charlesworth, 2018). The second mutation usually is dominant and causes female sterility resulting in a dioecious population of staminate individuals and pistillate individuals (Charlesworth, 2018). This transition from hermaphroditism to dioecy may have been driven by a reallocation of reproductive resources, which improves overall pollen and seed production (Charlesworth and Charlesworth, 1978).

While this transition may be beneficial, it is not necessarily unidirectional, even in systems with sex chromosomes. Reverse sexual transitions may be caused by environmental factors such as hormones and temperature (Ming et al., 2007; Ogata et al., 2016) or if the sexual strategy is advantageous for the population (Ming et al., 2011). The reverse sexual transition of males in a dioecious population to hermaphrodites results in a gynodioecious population. Reversions have been found in species where population bottlenecks occurred, due to shifts in population size and subsequent loss of genetic diversity (VanBuren et al., 2015). Also, major changes associated with polyploidy such as chromosomal rearrangements, changes in DNA sequence, and gene expression levels, may drive shifts in sexual systems, including reversions (Chen, 2007; Glick et al., 2016). While advantageous, this reverse sexual transition has only been studied in a few species, including *Datisca glomerata* (Durango root), *Mercurialis annua* (annual mercury; (Pannell et al., 2008), and *Carica papaya* (papaya).

Papaya is in the Caricaceae family, which consists of 35 different species, of which 32 are dioecious, two are trioecious and one is monoecious (Ming et al., 2007). Wild populations of papaya are dioecious while cultivated populations are generally gynodioecious (Brown et al., 2012). Sex in papaya is determined by sex chromosomes approximately 7 Ma (Wang et al., 2012), and papaya has served as a model to study the early stages of sex chromosome evolution (Wang et al., 2012; Weingartner and Moore, 2012; Wu and Moore, 2015). Papaya females are homogametic (XX), males are heterogametic (XY), and hermaphrodites are heterogametic with an alternative Y chromosome called the Y^h^ chromosome (XY^h^; Aryal and Ming, 2014). The Y and Y^h^ have non-recombining <u>s</u>ex-<u>d</u>etermining <u>r</u>egions (SDRs) called the <u>m</u>ale <u>s</u>pecific region of the Y (MSY) and the <u>h</u>ermaphroditic <u>s</u>pecific region of the Y^h^ (HSY), respectively.

The emergence of the hermaphroditic Y^h^ caused a reversion from dioecy to gynodioecy approximately 4,000 yrs ago (VanBuren et al., 2015). In natural papaya populations, there are three MSY haplotypes: MSY1, MSY2, and MSY3 (Weingartner and Moore, 2012; VanBuren et al., 2015). When compared to the three haplotypes, the HSY is most similar to MSY3, with fewer single nucleotide polymorphisms (VanBuren et al., 2015). Because of this similarity, the HSY most likely diverged from MSY3 haplotype.

The MSY3 and HSY are approximately the same size (~8Mb; Wang et al., 2012) and are gene poor relative to the pseudoautosomal region (PAR) of the chromosome (VanBuren et al., 2015). The MSY3 and HSY contain 94 total transcriptional units (TU), with 66 protein-coding genes and 28 pseudogenes (Wang et al., 2012). Both the MSY3 and HSY are highly repetitive and are flanked by PARs, which recombine like autosomes (Wang et al., 2012; VanBuren et al., 2015). The MSY3 and HSY are also between 98.8-99.6% identical in protein-coding sequences, with overall conserved gene content, gene order and exon structure (Wang et al., 2012; VanBuren et al., 2015). While similar genetically, male and hermaphroditic flowers have major morphological differences. Male staminate flowers have a pistillode, which is a sterile pistil that aborts early in development while hermaphroditic flowers have pistils that fully develop (Ronse Decraene and Smets, 1999).

Because the male Y is predicted to have a dominant gynoecium-suppressing (GS) locus, it is reasonable to hypothesize that this locus is non-functional in the Y^h^. Furthermore, because there are no significant coding differences in MSY and HSY protein-coding genes, it is likely that expression of the dominant GS locus in the HSY is impaired. Differences in male and hermaphrodite gene expression early in floral development, prior to or during the formation of the pistillode, are likely significant.

Therefore, we predict that the morphological differences between males and hermaphrodites as well as the cause of the reverse sexual transition in papaya is due to differentially expressed genes (DEGs), in particular the GS gene. To test this prediction, we performed RNA-seq of early and late developmental floral buds of males, females and hermaphrodites. We compared gene expression of X, Y and Y^h^ linked genes during floral development. We found sex-biased DEGs distributed across the sex chromosomes, with a substantial proportion in the PARs. Total gene expression of SDR-linked genes in the oldest evolutionary strata was roughly equal between males and females, and X-linked expression in XX females was roughly double that of X-linked expression in XY males and XY^h^ hermaphrodites in the two oldest evolutionary strata of the SDR. This suggests dosage compensation, a process which equalizes male and female X-linked expression, has not evolved for most sex-linked DEGs. In contrast, an analysis of hemizygous X-linked loci found evidence of partial dosage compensation. Finally, we identified a candidate sex determining gene that may be associated with the transition to hermaphroditism.

## MATERIALS AND METHODS

### Plant Material

Male (XY) and female (XX) wild papaya individuals were derived from a single maternal family in the Nicoya peninsula of Costa Rica (CR85 from Brown et al., 2012). All wild males in the study have the MSY3 haplotype, from which the HSY is derived (VanBuren et al., 2015). Hermaphrodites originated from the Kaek Dum Cultivar (XY^h^; HCAR3O9 from the Hawaii Agricultural Research Station, U.S. National Plant Germplasm System). All plants were grown in the Belk Greenhouse at Miami University.

Early and late developmental floral buds from five males, three females and three hermaphrodites (21 total samples) were collected in eppendorf tubes, flash-frozen in liquid nitrogen, and stored at −80C. Early developmental, pre-meiotic buds were 1-2 mm in length and collected prior to pistillode initiation in males and pistil initiation in females and hermaphrodites. Late developmental buds were 1 cm in length, in pre-anthesis (before petals opened), and were collected after pistillode initiation and subsequent abortion in males and after pistil formation and growth in females and hermaphrodites (Appendix S1). Male anthers in late developmental flowers were fully formed and contained mature pollen grains. RNA was extracted from one late developmental bud (one late developmental sample) and five early developmental buds (one early developmental sample) per individual using the Maxwell 16 LEV Plant RNA Extraction Kit (Promega Corporation, Madison, WI).

Total RNA quality was assessed using the Agilent 2100 Bioanalyzer (Agilent Technologies, Inc., Santa Clara, CA), and RNA Integrity Numbers (RINs) ranged between 7-9.5 (Schroeder et al., 2006). KAPA Stranded mRNA-Seq kits (Kapa Biosystems, Willmington, MA) with the Illumina platform were used to construct libraries sequenced by NovoGene, Inc. (Chula Vista, CA). Quality assessment of each library was performed using a Qubit 2.0 (ThermoFisher Scientific, Waltham, MA), an Agilent 2100 bioanalyzer (Agilent Technologies, Inc., Santa Clara, CA), and qPCR. Libraries were sequenced using paired-end 150 bp Illumina reads. Raw sequences were trimmed using Trimmomatic version 0.36 (Bolger et al., 2014) to remove adapter sequences and low quality leading and trailing regions (value 3). Trimmomatic scanned and cut reads with low average quality (value 15 across average four bases) and retained reads with a minimum length below 36 base pairs. The quality of trimmed reads was assessed using FastQC (Andrews). All raw RNA-seq reads are deposited in NCBI BioProject PRJNA640355.

Hierarchical Indexing for Spliced Alignment of Transcripts 2 (HISAT2; Pertea et al., 2016) was used to map the clean sequence reads to the papaya genome (Retrieved from: lftp://ftp.ncbi.nih.gov/genomes/Carica_papaya/; Ming et al., 2008), and previously assembled X- and HSY-sequenced regions (Wang et al., 2012). StringTie (1.3.4c) assembled mapped alignments and quantified transcript abundance for each library with default parameters (Pertea et al., 2016). Each library’s alignments were run through StringTie individually, and assembled transcripts were merged using “stringtie merge” to produce a unified annotation. The merged annotation file was processed into a matrix file using R (R Core Team, 2013).

Differential expression among six comparisons was measured using DESeq2 (Love et al., 2014). These comparisons included male versus female, male versus hermaphrodite, and female versus hermaphrodite at both early and late stages of development. Independent filtering of DEGs was conducted based on the mean of normalized counts (Love et al., 2014). All samples were analyzed using a Principal component analysis (PCA) in R (Appendix S2). DEGs with a significant *P* value (< 0.05) were extracted and annotated using blastx (Altschul et al., 1990) against the UniprotKB database (Chen et al., 2017).

Physical locations of the DEGs were identified using the papaya reference genome (Accession number: GCF_000150535.2), with the supercontigs listed by NCBI (Ming et al., 2008). Supercontigs were mapped to 12 linkage groups in papaya (Yu et al., 2009), with linkage group 1 corresponding to the sex chromosomes. We used the order of supercontigs along LG1, coupled with the bp count of each supercontig to obtain minimum physical distances of supercontigs along the sex chromosomes. DEGs were also mapped to previously sequenced X- and HSY-specific regions of the sex chromosomes (Wang et al., 2012).

## RESULTS

### mRNA sequencing and differential expression

RNA-seq libraries yielded approximately 7,000,000 to 42,000,000 raw paired reads which, following removal of low-quality sequences, resulted in in a range of 6,000,000 to 39,000,000 cleaned paired reads (Appendix S3). The percent alignment with the reference genome ranged from 77%-84%. Over 95% of all samples had a Q20 score, and between 90-92% of all samples had a Q30 score (Cock et al., 2010). Overall, the number of paired cleaned reads and percent alignment were acceptable quality in order to perform differential expression analyses (Love et al., 2014).

Six pairwise comparisons of gene expression were performed to identify DEGs involved in reproductive differentiation among male, female, and hermaphroditic sexes at early and late developmental stages using DESeq2. We refer to genes that exhibit differential expression between two floral sexual types as sex-biased DEGs, including differences between hermaphrodites and either males or females. The comparison between late males versus late females had the most DEGs (3130), followed by the comparison between late males and late hermaphrodites (1417). In a PCA of variability among samples, late males were distributed widely across PC1 and PC2, whereas the rest of the samples are more tightly clustered along PC1 with the exception of early male sample 6 (EM6; Appendix S2).

### Genomic distribution of differentially expressed genes

We identified a total of 4145 DEGs from the six developmental comparisons among the sexes representing 3021 unique transcripts, of which 217 localized to linkage group 1 (LG1), previously identified as the sex chromosome (Yu et al., 2009). Of the other unique transcripts, 1644 localized to the autosomes (LG2-12; Yu et al., 2009), while 1054 were found on supercontigs not mapped to linkage groups. Of the remaining loci, 92 mapped to the X and/or HSY (see below), while 25 mapped to the chloroplast genome.

Of the DEGs found in LG1, 21 localized to supercontigs previously identified as spanning the SDR (supercontigs 39, 315, 794, and 66, Appendix S4); therefore, the majority (89.8%) of the DEGs on LG1 localized to the PARs (Fig. 1). However, the proportion of PAR-linked DEGs was not significantly different than that for the rest of the genome (Appendix S5; *P* > 0.05, Fisher’s Exact Test). A simple linear regression of the number of DEGs per sex chromosome supercontig versus the physical distance of that supercontig to the SDR showed no significant correlation between DE gene density and physical distance to the SDR (*r^2^* = 0.0191, *P* > 0.05; Appendix S6).

**Figure 1.**
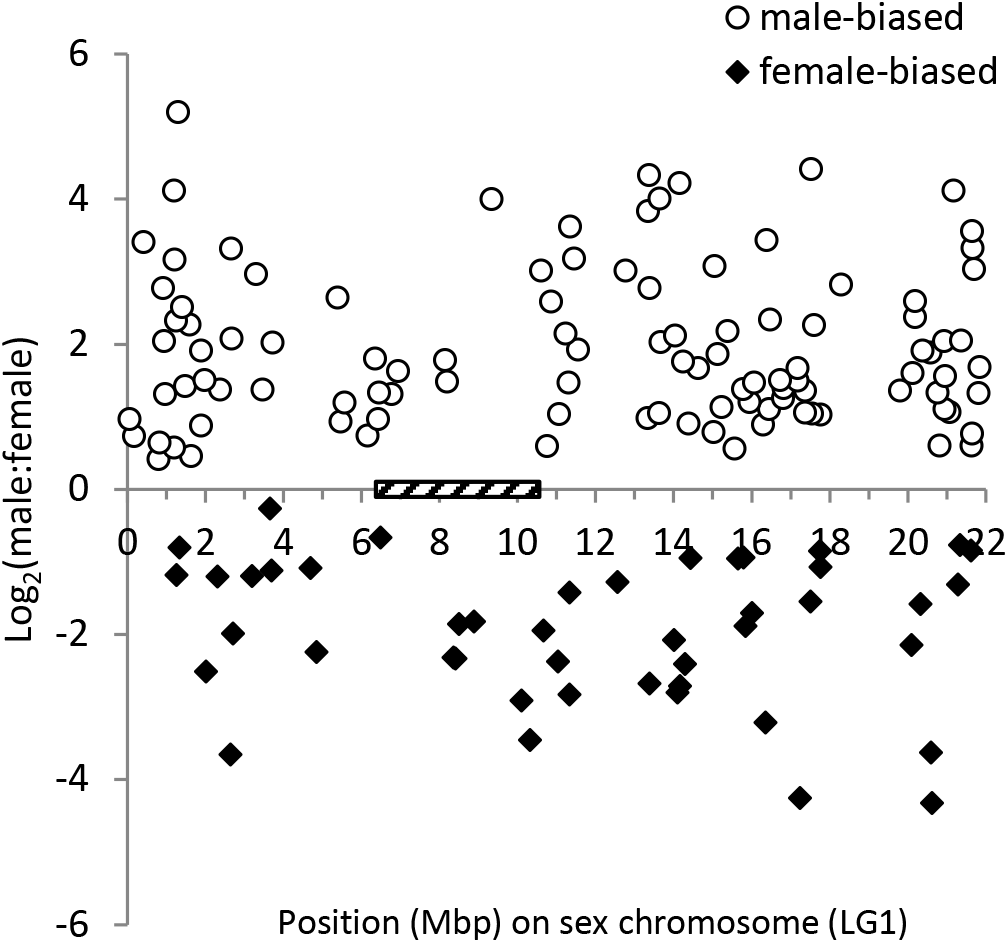
Log_2_ ratio of male to female expression for male-biased and female-biased differentially expressed genes (DEGs) versus position on the sex chromosome (Linkage Group1). The position of the sex-determining region (SDR) is indicated by the boxed region.

An analysis of sex-biased expression of LG1 DEGs found there is a significant excess of male-biased DEGs (108, significantly upregulated in males versus females) relative to female-biased DEGs (46, significantly upregulated in females versus males) localized to the sex chromosomes (P < 0.001, Fisher’s exact test; Fig. 1). In addition, 71 DEGs were significantly upregulated in males relative to hermaphrodites, 73 DEGs were significantly upregulated in females relative to hermaphrodites, and 8 DEGs upregulated in both males and females relative to hermaphrodites.

### Differentially expressed genes in the X- and HSY sequenced regions

We identified 109 DEGs mapping to the X- and HSY sequenced-regions representing 92 loci (Table 1). The SDR of the X-sequenced regions is preceded by an ~ 1.9 Mb flanking PAR region (Fig 2). DEGs in the X-PAR showed no clear pattern of sex-biased expression, with equal numbers of male- and female-biased loci (Appendix S7). However, within the SDR, 23 of 25 X-linked loci were female-biased and 31 of 32 Y-linked loci are male-biased; most of these loci were X/Y gametologs (Appendix S8).

**Table 1.** Number and type of sex-biased DEGs in the PAR and SDR of the X- and HSY-sequenced regions.

| Expression Bias | X-PAR | Y-PAR | SDR |  |  |  |
| --- | --- | --- | --- | --- | --- | --- |
|  |  |  | X-only | Y-only | XY-X | XY-Y |
| Male/Female sex-biased |  |  |  |  |  |  |
| Male | 7 | 0 | 1 | 12 | 2 | 19 |
| Female | 7 | 0 | 4 | 0 | 18 | 1 |
| Other sex-biased |  |  |  |  |  |  |
| Male vs. Hermaphrodite |  |  |  |  |  |  |
| Male-biased | 5 | 0 | 1 | 7 | 0 | 0 |
| Herm-biased | 1 | 0 | 0 | 0 | 0 | 0 |
| Female vs. |  |  |  |  |  |  |
| Hermaphrodite |  |  |  |  |  |  |
| Female-biased | 2 | 8 | 0 | 1 | 0 | 0 |
| Herm-biased | 6 | 8 | 0 | 2 | 0 | 0 |
| Total | 28 | 16 | 6 | 22 | 20 | 20 |

**Figure 2.**
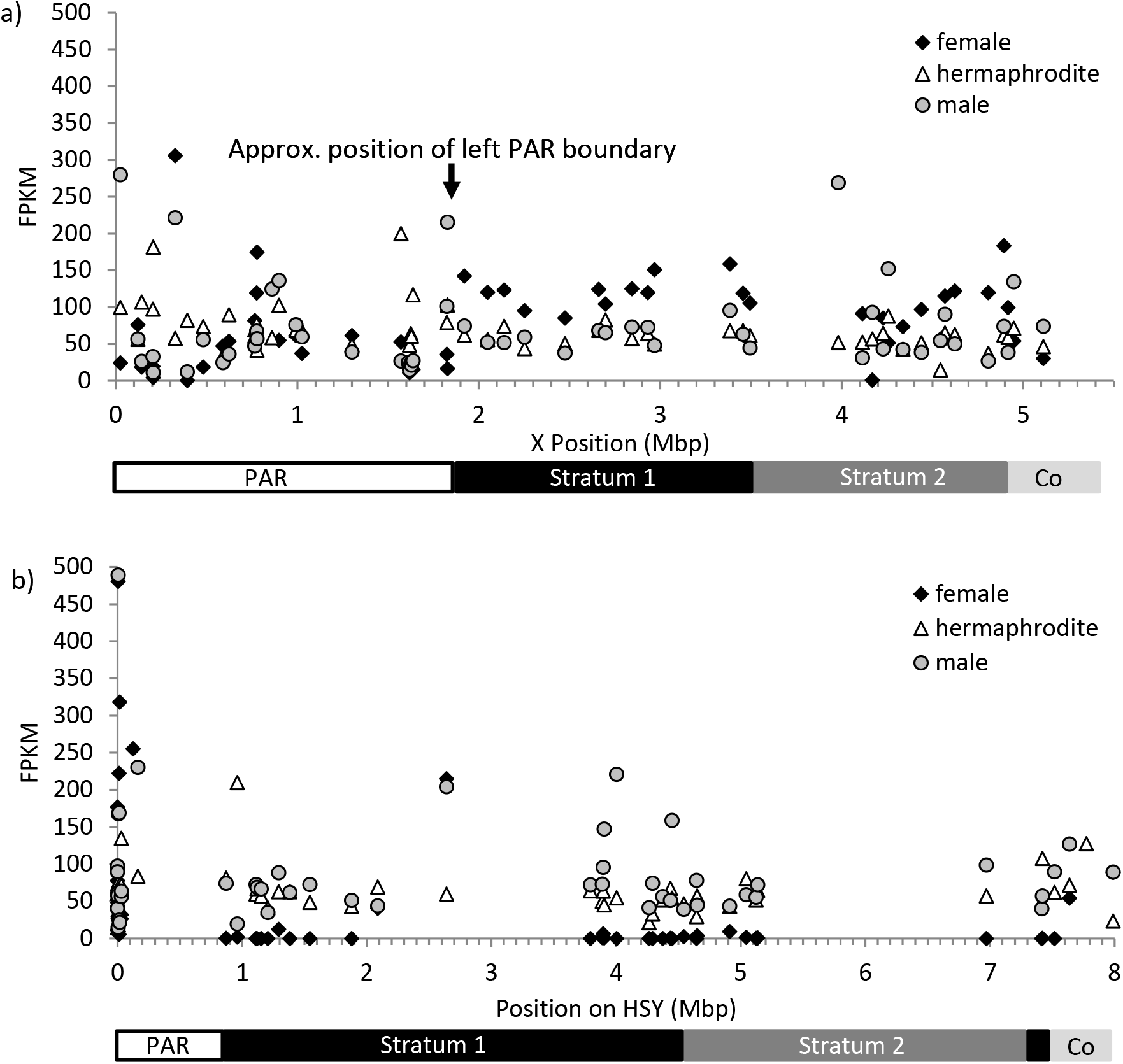
Total expression (FPKM) of differentially expressed genes (DEGs) in female (diamond), hermaphrodite (triangle), and male (circle) early- and late-developmental floral stages versus position along the sequenced (a) X-specific region and (b) hermaphroditic specific region of the Y (HSY). Approximate locations of the pseudoautosomal region (PAR), Stratum 1, Stratum 2, and the Collinear (Co) in the sequenced X and HSY sequenced regions is indicated below each graph.

Other sex-biased DEGs showed expression differences between either males or females and hermaphrodites. When comparing males and hermaphrodites, most of X-linked DEGs were found in the X-PAR (6 of 7 loci), and the majority of these loci showed up-regulation in hermaphrodites (5 of 7 loci; Table 2). In contrast, all 7 of the Y-linked DEGs when comparing males and hermaphrodites were upregulated in males (Table 2). When comparing females and hermaphrodites, all of the DEGs show differential expression specifically in late developmental tissue (Appendix S9). Within the X-PAR, 6 DEGs showed upregulation in hermaphrodites, while 2 DEGs were upregulated in females. Two of the three Y-linked DEGs were upregulated in hermaphrodites relative to females (Appendix S9).

**Table 2.**
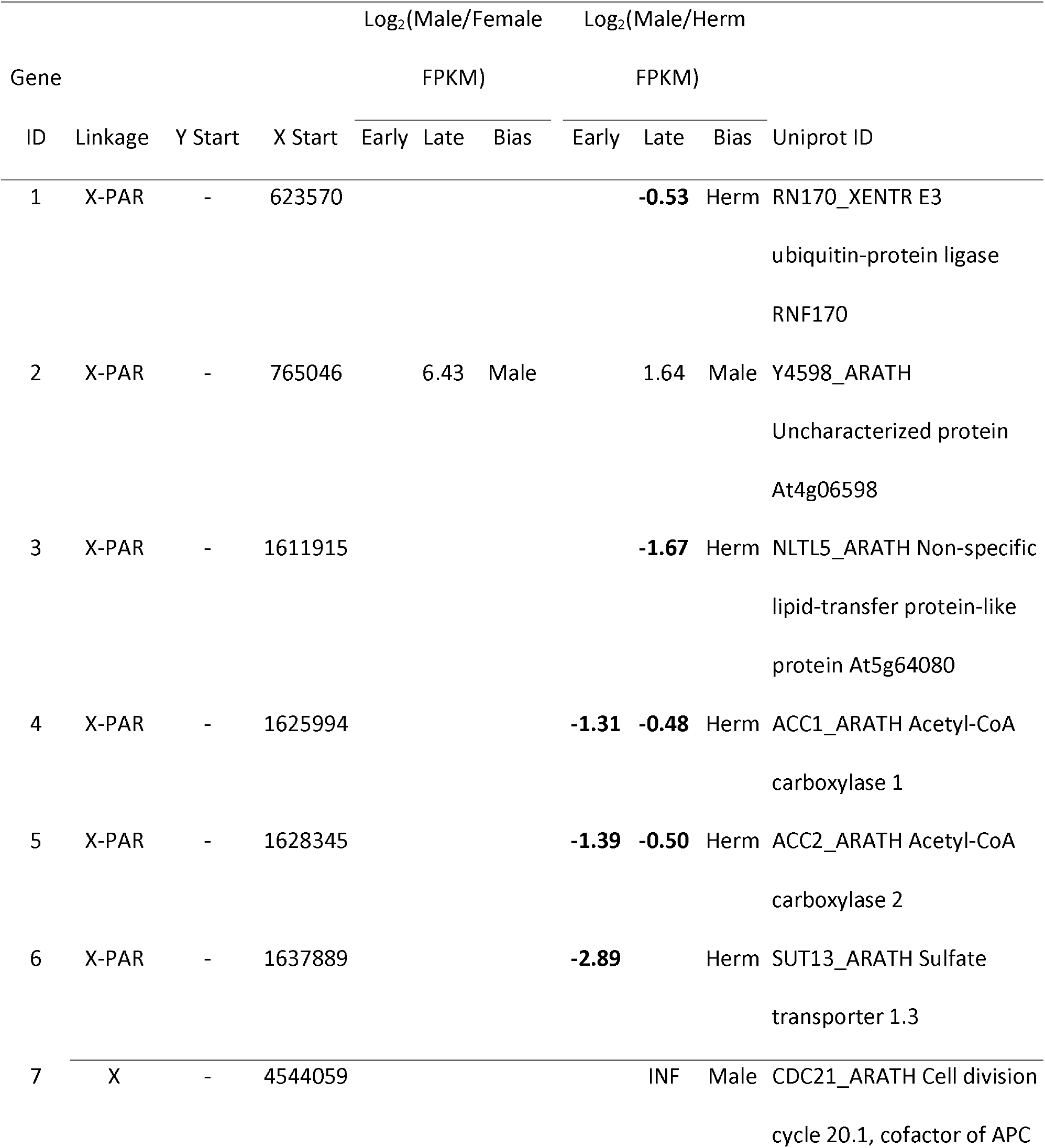
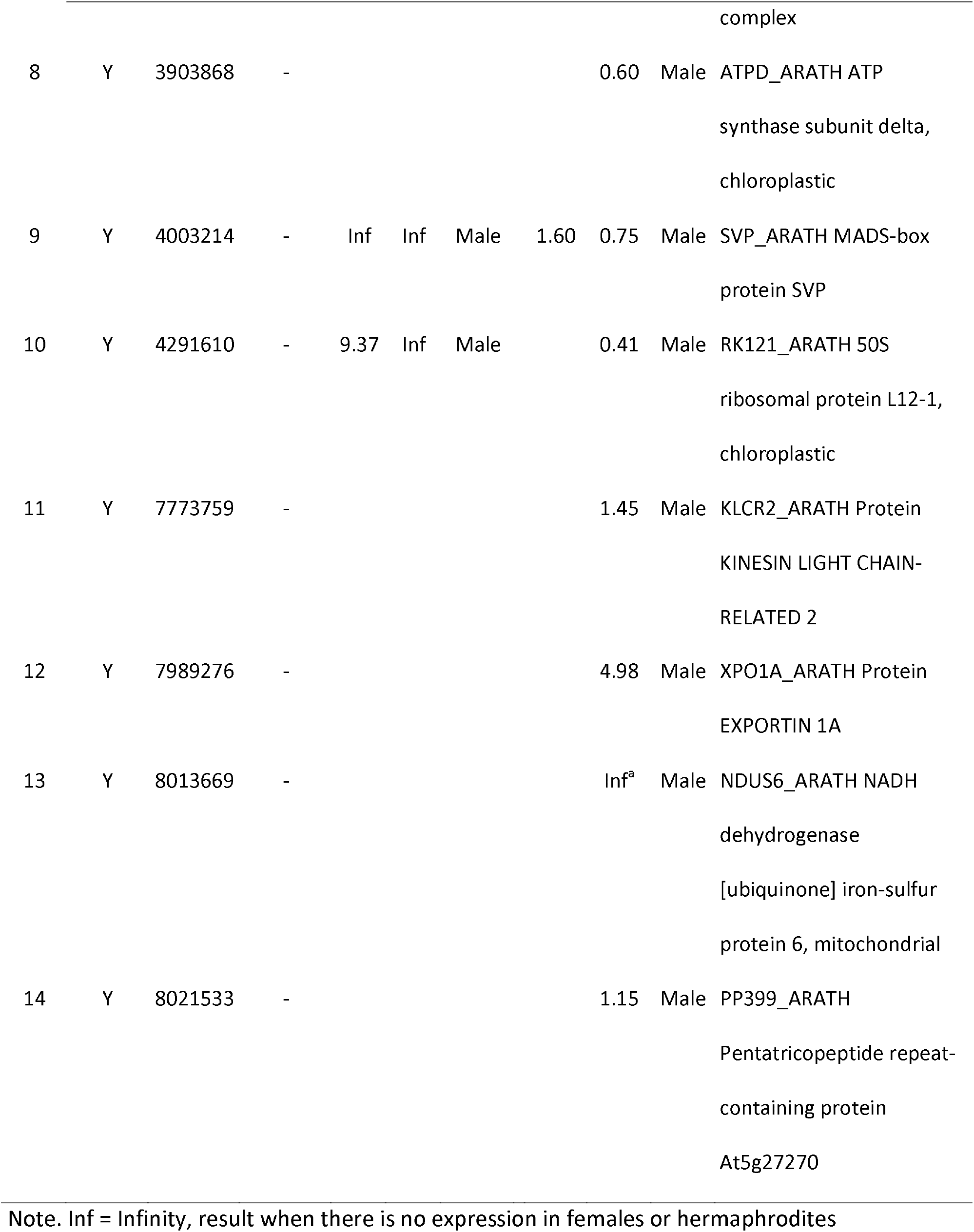
Location and expression bias of DEGs mapping to the X- and HSY-sequenced regions comparing male versus female expression and male versus hermaphrodite early and late floral development.

We investigated these patterns of expression more closely in the 20 DEG loci with X- and Y-gametologs for evidence of dosage compensation; we also included an additional 20 X/Y gametolog pairs for which we detected expression, for a total of X/Y gametolog pairs. If dosage compensation is occurring in the papaya SDR, we expect to equivalent expression of X-linked loci in females (with two X-copies) and X-linked loci in males (with one X-copy). If dosage compensation is not occurring, we expect twice as much expression of X-linked loci in females relative to the X-linked gametolog in males; furthermore, the sum of X-and Y-linked gametolog expression in males will be equivalent to X-linked expression in females. We divided the analysis according to strata, with 14 X/Y pairs in the oldest evolutionary stratum 1, 16 X/Y pairs in stratum 2, and 10 X/Y pairs in the youngest, collinear region (Appendix S10).

The average ratio of X-linked expression in males to XX females (X male: XX female) was 0.56 ± 0.04 SE in stratum 1, 0.52 ± 0.08 SE in stratum 2, and 1.0 ± 0.2 SE for the collinear region (Fig 3). Combined expression of X and Y gametologs in males was roughly equal to that found for total expression of the two X alleles in females for the majority of genes in the first two evolutionary strata, with an average ratio of total male expression to total female expression of 1.1± 0.08 SE in stratum 1, and 1.1 ± 0.1 SE in stratum 2 (Fig. 4, Appendix S10). One locus in stratum 2 has at least twice the total expression in males relative to females (gene X/Y pair 9, Fig 4). In the collinear region, the ratio of total male expression to total female expression was much higher, with an average value of 2.4 ± 0.5 SE (Fig. 4).

**Figure 3.**
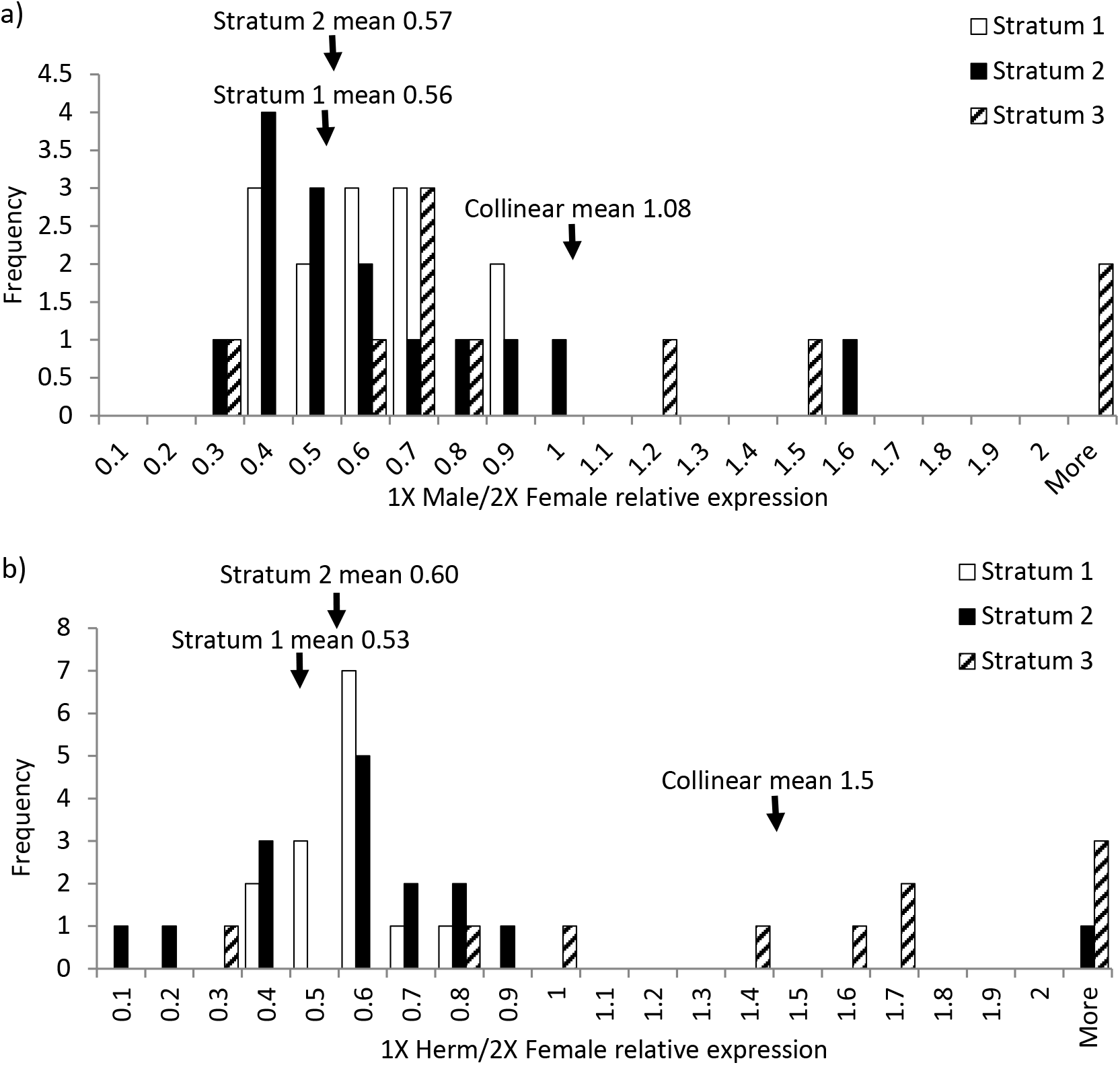
Frequency distribution of (a) male and (b) hermaphrodite X-linked expression relative to 2X female expression for 40 SDR-linked genes with X- and Y-linked gametologs. Expression data is divided across the three different evolutionary strata of the sex-determining region (SDR), Stratum 1 (white), Stratum 2 (black), and the collinear region (striped).

**Figure 4.**
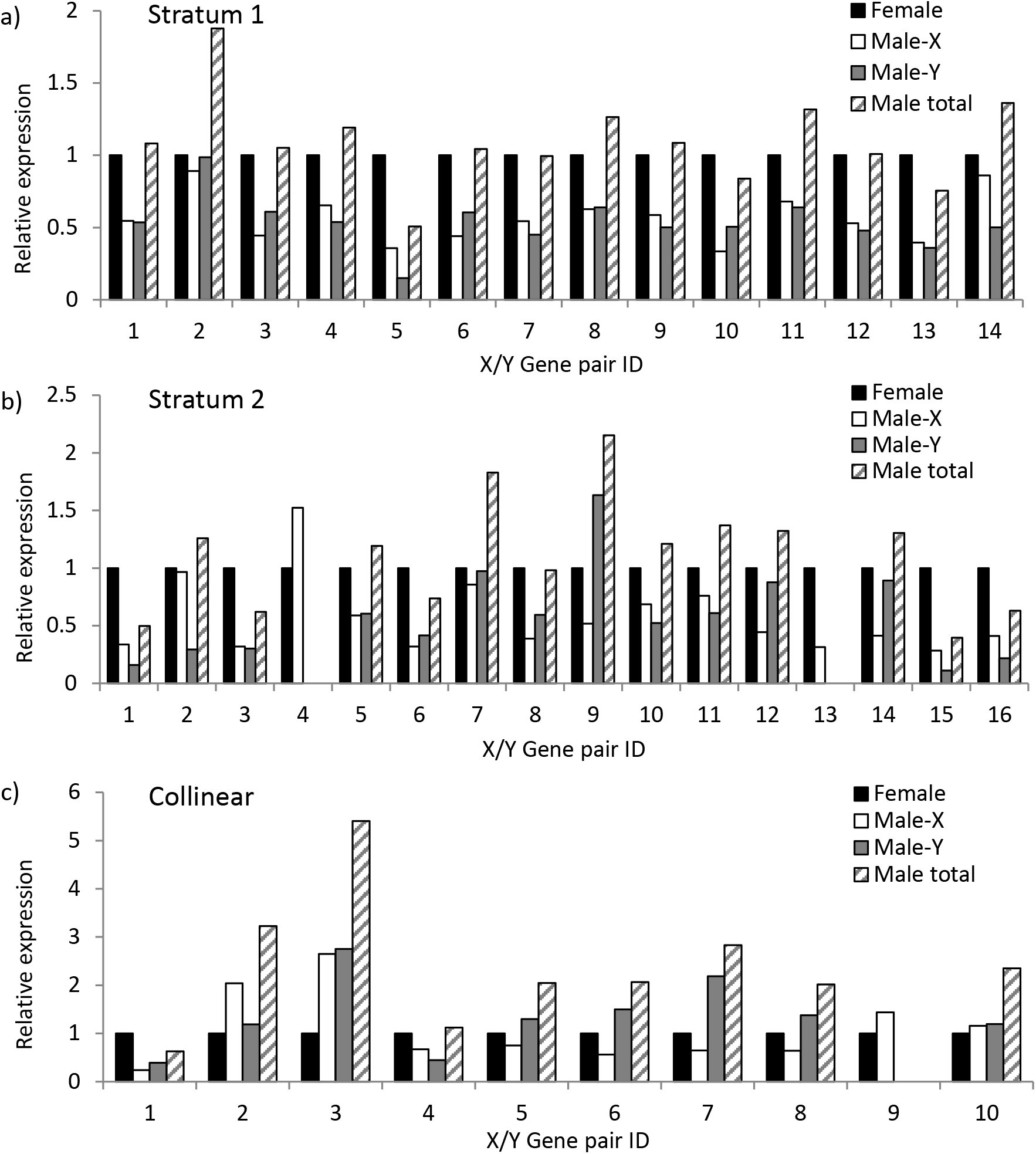
Expression of male X- and Y-linked gametologs and total male expression relative to total female expression for 40 sex-determining region (SDR) genes with X- and Y-linked gametologs. Data for each gene is presented according to the evolutionary stratum in which they are found: (a) Stratum 1, (b) Stratum 2, (c) Collinear region. Gene pair IDs are described in Appendix S10.

It has been shown in other young plant sex chromosome systems that dosage compensation may be limited to X-linked gametologs with degenerate or non-expressed Y gametologs (Muyle et al., 2012; Papadopulos et al., 2015.) In order to test this possibility in papaya, we identified 19 hemizygous X-linked genes; i.e., they lack a Y-gametolog with observable expression (11 genes from stratum 1, four genes from stratum 2, and three genes from the collinear region). The average ratio of total expression for these hemizygous genes in males (X) to females (XX) was 0.86, with some loci having close to equivalent expression (Fig. 5). However, the distribution of male:female (M:F) expression was broad and bimodal, ranging from 0.30 to 1.9 (Fig. 5). Furthermore, mean male to female expression was lower in stratum 1 and 2 (mean M:F of 0.78 and 0.72 respectively) than in the collinear region (mean M:F of 1.3).

**Fig. 5.**
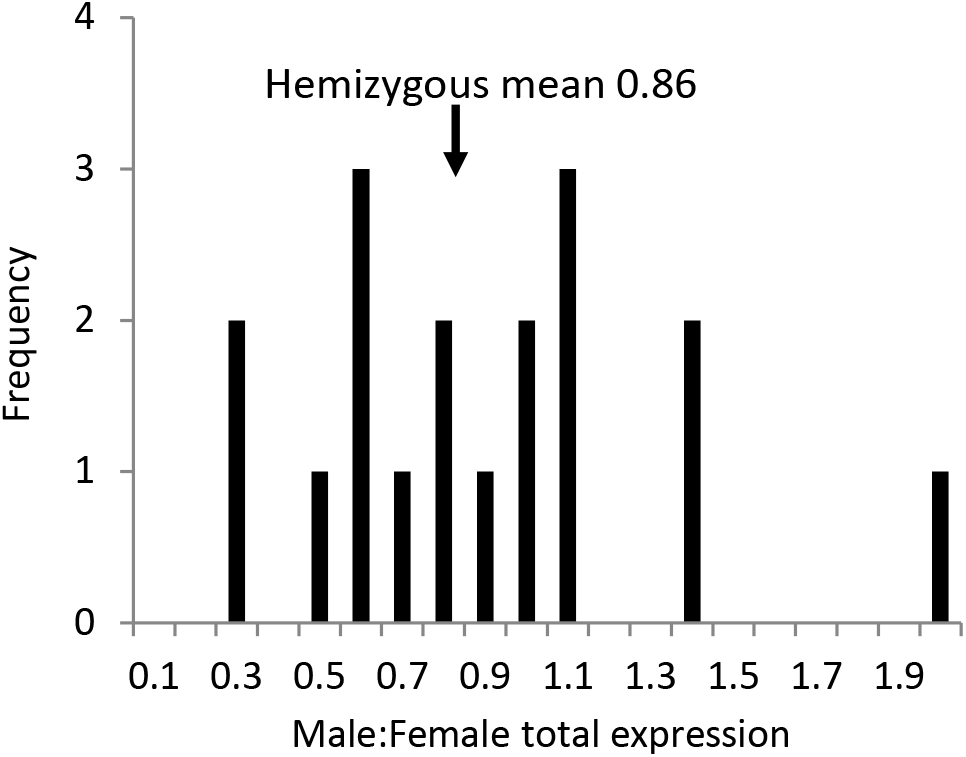
Frequency distribution of total male to female expression of 19 hemizygous genes.

### Candidate genes involved in the male-to-hermaphrodite transition

The transition from male to hermaphrodites predictably involves the loss-of-function of a dominant gynoecium suppressing locus on the Y chromosome. We predict that such a locus should be up-regulated in male development, resulting in gynoecium abortion, and down-regulated in hermaphrodites and females. There were 14 DEGs that had significantly different expression patterns between males and hermaphrodites (Table 2), of which seven were Y-linked. All seven were significantly up-regulated in males relative to hermaphrodites; however, five were specific to only the late-stage flowers, and were not significantly up-regulated in male flowers relative to female flowers. The remaining two loci show up-regulation in male flowers versus both female and hermaphrodites; however, only one of these is significantly up-regulated in both early and late developmental stages, a homolog of the MADS-box protein *SHORT VEGETATIVE PHASE* (*SVP*) from *Arabidopsis thaliana* (Table 2, Gene ID 9).

## Discussion

### Sex-linked differentially expressed floral developmental genes are found disproportionally on the PAR

Sex-biased expression is found in many species, though it may vary according to developmental stage, with low levels of sex-bias in embryonic stages to high sex-bias in sexually mature adults (Grath and Parsch, 2016). Sex-biased gene expression eliminates the negative effects of sexually antagonistic alleles in the sex where it is harmful (Connallon and Knowles, 2005; Scotti and Delph, 2006). To negate the harmful effects of sexually antagonistic alleles, species may evolve by developing linkages between sexually antagonistic mutations found in the PARs and the SDR (Vicoso et al., 2013). These linkages lead to this mutation being transmitted more to the sex that benefits more (Vicoso et al., 2013). Therefore, sex-biased genes are expected to be preferentially found not only in the SDR but also in the PARs (Otto et al., 2011), implying that genes beneficial to each sex are more likely to be on sex chromosomes compared to autosomes (Bachtrog et al., 2011).

Similar patterns are observed in both plants and animals. For example, PAR-linked sex-specific quantitative trait loci (QTLs) were identified on the sex chromosomes of the dioecious plant *Silene latifolia* (Delph et al., 2010). There were eight male sex-specific QTLs compared to one female sex-specific QTL in the PAR, which represents 10% of the sex chromosome (Delph et al., 2010). PAR-linked sex-biased expression has also been observed on the ZW sex chromosomes of ratite birds, such as emus and ostriches (Vicoso et al., 2013). The W-linked SDR of ratite birds is also restricted in size, resulting in PARs approximately two-thirds the size of the sex chromosome (Yazdi and Ellegren, 2014). There is substantial PAR-linked sex-biased expression in ratite sex chromosomes (Vicoso et al., 2013). When gonads form, male-biased gene expression on the Z, including the PARs, increases, suggesting that the Z has become masculinized (Vicoso et al., 2013).

In papaya, most of sex-biased DEGs on the sex chromosomes were evenly distributed across the sex chromosomes, with the majority found in the larger PARs. However, we did not find an overabundance of sex-biased DEGs on the sex chromosomes; indeed, there were proportionally similar frequencies of sex-biased DEGs locating to the autosomes. And while it is predicted for PAR-linked sexually antagonistic genes to accumulate nearer the SDR (Otto et al., 2011; Qiu et al., 2013; Charlesworth et al., 2014), we found that sex-biased gene expression was distributed evenly across the papaya PARs. It is possible that sexualization of sex chromosomes may take longer to achieve, as papaya sex chromosomes evolved roughly 7 Ma, whereas the sexualized sex chromosomes of ratite birds are approximately 120 Ma (van Tuinen and Hedges, 2001). Our results do further emphasize the critical role that the PAR may play in sex chromosome evolution, and further investigations into the role PAR-linked genes play in sexual differentiation could shed light on this phenomenon (Otto et al., 2011).

### SDR-linked DEGs and sex-biased gene expression

Sex-biased gene expression was found in both the PAR and SDR regions of the X- and HSY-sequenced regions; however, the PAR-linked genes showed equivalent numbers of male- and female-biased DEGs. In the SDR, there were 37 loci with male/female sex-biased expression; almost all X-linked loci are female-biased, while Y-linked loci are male-biased. However, 20 of these loci are single gene loci with X- and Y-gametologs, and while X-linked gametologs are female-biased and Y-linked gametologs are male-biased, this is more consistent with gene dosage effects. X-linked gene expression in females for 18 of these 20 loci is, on average, twice that of X-linked gene expression in males. Overall expression of these genes in males, when summed across X- and Y-linked expression is equal to that of females, thus, these loci are not sex-biased.

In addition, when we survey gene expression across the SDR, including all genes with expressed X- and Y-gametologs, we found no clear signature of dosage compensation. In the oldest evolutionary strata 1 and 2 (14 and 16 loci respectfully), total male and female expression was approximately equivalent, while X-linked male expression was roughly one-half that of XX females. Studies of dosage compensation in the young plant sex chromosomes of S. *latifolia* suggest that partial dosage compensation does exist in that system, though the signature of dosage compensation is strongest in sex-linked genes with degenerated or non-expressed Y-linked gametologs (Muyle et al., 2012; Papadopulos et al., 2015; Muyle et al., 2018; Krasovec et al., 2019). We also found evidence of dosage compensation in X-linked hemizygous loci. As in previous studies, the pattern of DC was partial, with average male to female expression exhibiting a bimodal distribution (Muyle et al., 2012; Papadopulos et al., 2015). Furthermore, the signature of dosage compensation was distributed among evolutionary strata, suggesting dosage compensation may be evolving on a gene-by-gene basis.

The story is more complex for collinear genes. The collinear region is adjacent to stratum 2, and is not found in a large-scale chromosomal inversion (Wang et al., 2012). It is characterized by low divergence rates between X and Y gametologs, and some loci have shared polymorphism (Wang et al., 2012; Wu and Moore, 2015). Unlike Stratum 1 and 2, the 10 loci in the collinear region had greater total expression in males than in females and equivalent X-linked expression in males relative to XX females. These results are consistent with dosage compensation in this region; however, low divergence between X and Y gametologs could confound assignment of transcripts to the X and HSY (i.e. the same transcript may map to both regions, leading to a false signature of dosage compensation). Thus, we suspect this result is an artifact of the transcript assembly process and not an indicator of dosage compensation.

### Investigating the male to hermaphrodite transition

The two-gene model of plant sex chromosome evolution predicts the presence of two sex-determining loci on the male Y chromosome; one is a dominant male-promoting locus and one a dominant gynoecium suppressing (GS) locus (Harkess et al., 2017; Charlesworth, 2018; Akagi et al., 2019; Harkess et al., 2020). The emergence of hermaphrodites in papaya supports this two-gene model, in that it represents an independent reversion, or loss-of-function mutation, of the gynoecium suppressing locus. Molecular dating supports the recent emergence of the HSY from the MSY approximately 4000 yrs ago, coincident with papaya domestication (VanBuren et al., 2015). As the MSY and HSY are nearly identical in protein-coding sequences (VanBuren et al., 2015), we predicted that the reversion to hermaphrodites involves a loss of gene expression in a GS gene on the HSY. This gene predictably acts in early male floral development, where it causes the gynoecium to abort and the formation of an underdeveloped pistillode.

Twelve DEGs were identified in comparisons of male and hermaphroditic developmental tissues (Table 2). Of the five Y-linked DEGs, all five were up-regulated in late male-development, but only one was also up-regulated in early male-development relative to hermaphrodites, MADS-box protein *SVP*, where it has a 9.9 log_2_-fold increased expression. This locus is also up-regulated in all male vs. female comparisons, with no expression detected in females. In addition, the SVP locus is the only one of the five HSY-linked candidates with a floral regulatory function (Gregis et al., 2006; Gregis et al., 2013). In other plant species, *SVP* represses floral transition and regulates B, C, and E class genes (Gregis et al., 2013). It may also play a role in the endodormancy of deciduous fruit tree species (Yamane et al., 2008).

It is possible that the GS may directly function in pistil abortion, leading to formation of the pistillode that is present in our late developmental floral stage. In a functional mapping study of papaya, the *SVP* gene had elevated expression in the third whorl of both male and female sterile flowers and low in the fourth whorl of hermaphrodites, consistent with a regulatory role affecting fourth whorl development (Lee et al., 2018). Genomic analyses confirm that the *SVP* allele is not present in the X chromosome, but is specific to the Y- and Y^h^ chromosomes (Wang et al., 2012; Lee et al., 2018). It is found in the oldest evolutionary stratum 1, which is where we would expect to find the original sex determining loci. Only the allele in the male Y chromosome has a copy of the *SVP* gene that encodes for an intact protein (Lee et al., 2018), implying that it may be associated with pistillode formation and abortion.

The specific role of *SVP* in pistillode formation, however, is unclear. First, *SVP* is primarily involved in the transition from the vegetative to floral development and overexpression of *SVP* in other systems, such as kiwi fruit, lead to a delay in flowering (Wu et al., 2012). Another study with barley also showed that *SVP* suppressed floral meristems and even caused flowers to lose floral identity and become an inflorescence (Trevaskis et al., 2007). While our analysis support SVP as a strong GS candidate locus, functional analyses of *SVP* need to be conducted in papaya to confirm its role in gynoecium development (Lee et al., 2018). It is also possible our analysis has missed a key regulator of gynoecium development if the GS is expressed earlier in development or at very low levels. As well, there could be that a specific microRNA (miRNA) or differential methylation contributed to the morphological reversion from males to hermaphrodites.

## Acknowledgements

The authors would like to thank our collaborators Ray Ming (University of Illinois at Urbana-Champaign) and Qingyi Yu (Texas A&M University) for providing cultivar seeds and sequence information. This work was supported by the National Science Foundation (NSF PGRP 1546890) as a subaward to RCM from the University of Illinois (PI, R. Ming).

## Author contributions

TC collected samples, performed molecular work, analyzed data, and wrote and edited the paper.

RCM assisted in data analyses, and wrote and edited the paper. AH assisted in computational analyses and edited the paper.

## Data availability Statement

All raw RNA-seq reads are deposited in NCBI BioProject PRJNA640355 (http://www.ncbi.nlm.nih.gov/bioproject/640355)

## Supporting Information

Additional Supporting Information may be found online in the supporting information section at the end of the article.

