## Supplemental Appendices for "Sex-linked gene expression and the reversion to hermaphroditism in *Carica papaya* L. (Caricaceae)"

List of supplemental tables

Appendix S1. Late floral developmental stages of females, hermaphrodites, and males.

Appendix S2. Principal component analysis plot showing variance among samples.

Appendix S3. Novogene QC summary of raw and cleaned paired end reads and percent alignment.

Appendix S4. LG1 supercontigs presented in order of physical position with NCBI location ID, the number of DEGs, and total genes per supercontig.

Appendix S5. Number of unique DEGs on LG1, the sex chromosome, and LG2-12, the autosomes, compared to the total number of genes in those regions.

Appendix S6. Relationship between DEG density (number of DEGs in supercontig/total number of genes in that supercontig) to distance of that supercontig from the SDR

Appendix S7. Location and expression bias of differentially expressed genes mapping to the X-PAR, X- and HSY-sequenced regions comparing male versus female early and late floral development.

Appendix S8. Location and expression bias of SDR-linked DEGs with X and Y gametologs comparing male versus female early and late floral development.

Appendix S9. Location and expression bias of DEGs with X-PAR-, Y-PAR-, and Y-linked DEGs comparing female versus hermaphrodite late floral development.

Appendix S10. Expression analysis male X- and Y-gametologs and total male expression relative to total female expression.

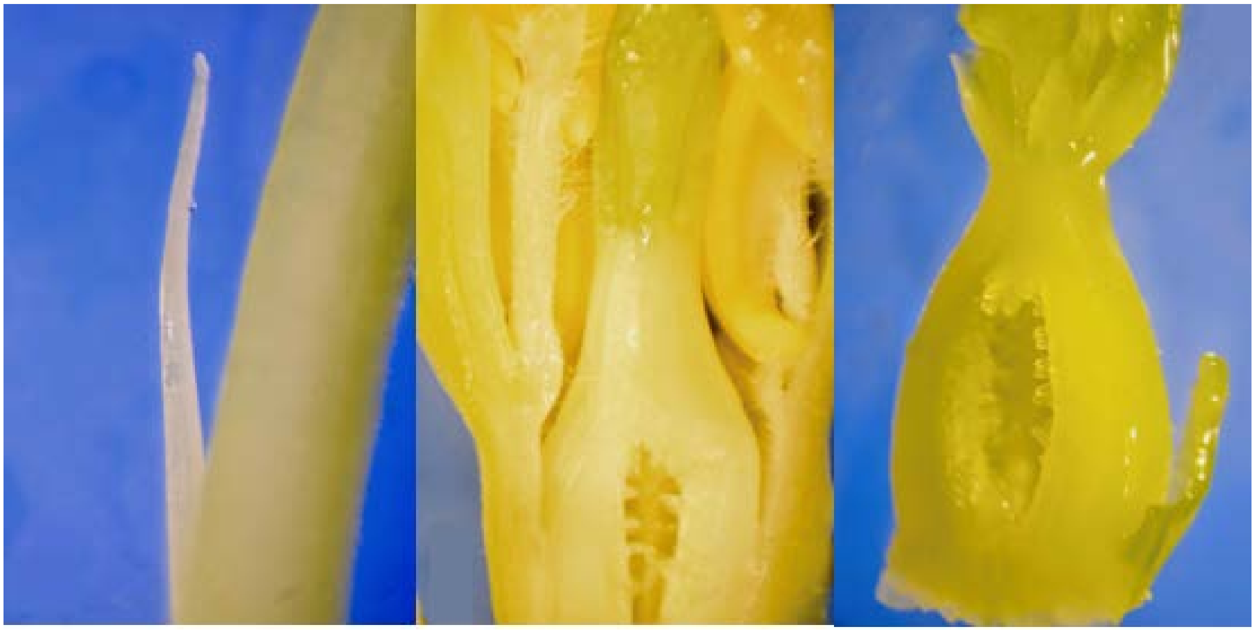

Appendix S1. Late floral developmental stages of females, hermaphrodites, and males. (A) Late developmental males (far right) have an aborted pistillode that looks like an undifferentiated gynoecium. (B) Mid to Late developmental hermaphrodites have an early forming gynoecium with a stigma and style that have not yet differentiated. (C) Female flowers in late development have a developed ovary, stigma and style.

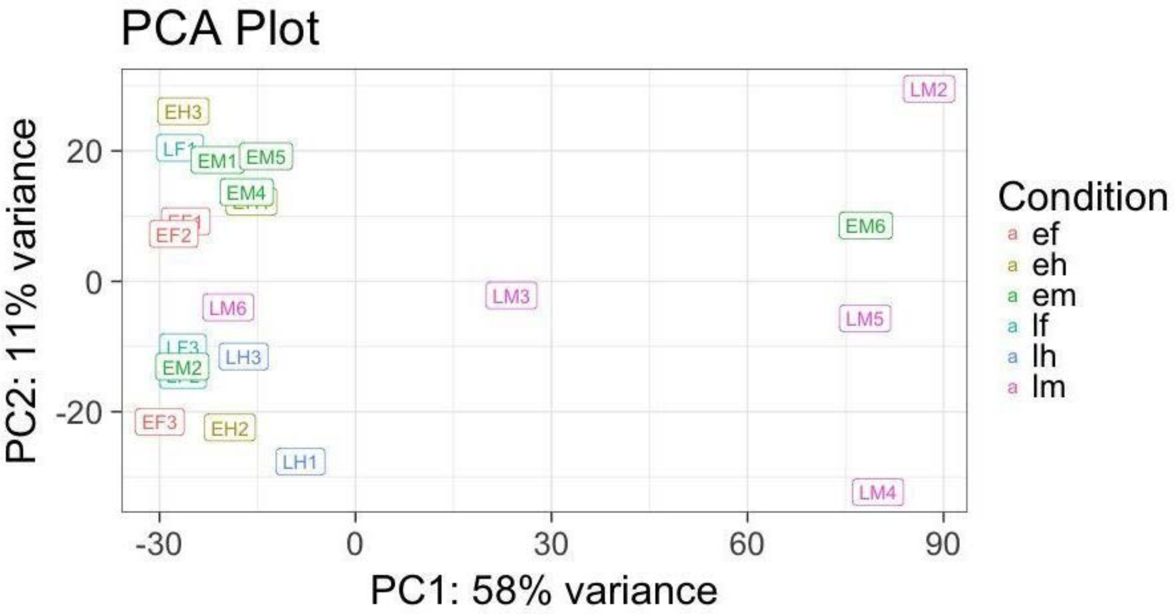


Appendix S2. Principal component analysis plot showing variance among samples. The first letter stands for either early (E) or late (L) development, while the second letter stands for either male (M), hermaphrodite (H), or female (F). The numbers stand for which tree from which the sample originated. For example, LH1 and EH1 are late hermaphrodite and early hermaphrodite samples respectively, and originated from the same individual.

Appendix S3. Novogene QC summary of raw and cleaned paired end reads and percent alignment

| Sample | Sample Sex/development | Raw paired reads | Cleaned paired reads | Percent Alignment |
|---|---|---|---|---|
| EH1 | Early hermaphrodite | 9222276 | 7379412 | 81% |
| EH2 | Early hermaphrodite | 14749039 | 11869558 | 79% |
| LH1 | Late hermaphrodite | 7220415 | 6026140 | 77% |
| LH2 | Late hermaphrodite | 18829895 | 15615380 | 79% |
| LH3 | Late hermaphrodite | 23266846 | 19331137 | 81% |
| LF1 | Late female | 12239268 | 9738847 | 84% |
| LF2 | Late female | 25199835 | 20247424 | 81% |
| LF3 | Late female | 22750274 | 17966252 | 81% |
| EF1 | Early female | 23299371 | 19255745 | 79% |
| EF2 | Early female | 26637013 | 21549664 | 82% |
| EF3 | Early female | 9865155 | 7958638 | 82% |
| LM2 | Late male | 7958638 | 16713003 | 84% |
| LM3 | Late male | 41054794 | 35945888 | 81% |
| LM4 | Late male | 19893127 | 17987837 | 78% |
| LM5 | Late male | 16871795 | 15489282 | 78% |
| LM6 | Late male | 17069405 | 15964421 | 78% |
| EM2 | Early male | 21800062 | 17282004 | 81% |
| EM1 | Early male | 42147140 | 39413395 | 83% |
| EM4 | Early male | 16690391 | 14539874 | 81% |
| EM5 | Early male | 17052433 | 15272980 | 80% |
| EM6 | Early male | 14608006 | 13471501 | 80% |

Appendix S4. LG1 Supercontigs presented in order of physical position with NCBI location ID, the number of DEGs, and total genes per supercontig.

| Supercontig Number | NCBI Location ID | Number of DEGs | Total Number of Genes |
|---|---|---|---|
| 116 | NW_019014576.1 | 11 | 93 |
| 1066 | NW_019013695.1 | 1 | 1 |
| 3603 | NW_019012029.1 | 0 | 1 |
| 14 | NW_019014678.1 | 50 | 261 |
| 140 | NW_019014552.1 | 9 | 54 |
| 99 | NW_019014593.1 | 8 | 64 |
| 123 | NW_019014569.1 | 5 | 60 |
| 39[a] | NW_019014653.1 | 18 | 112 |
| 215[a] | NW_019014477.1 | 3 | 11 |
| 794[a] | NW_019013934.1 | 0 | 3 |
| 66[a] | NW_019014626.1 | 19 | 115 |
| 177 | NW_019014515.1 | 5 | 24 |
| 160 | NW_019014532.1 | 12 | 32 |
| 2282 | NW_019012784.1 | 0 | 1 |
| 142 | NW_019014550.1 | 7 | 60 |
| 249 | NW_019014444.1 | 1 | 15 |
| 21 | NW_019014671.1 | 31 | 243 |
| 438 | NW_019014267.1 | 0 | 4 |
| 1076 | NW_019013688.1 | 0 | 1 |
| 36 | NW_019014656.1 | 35 | 207 |
| 49 | NW_019014643.1 | 22 | 161 |
| 1124 | NW_019013651.1 | 0 | 3 |
| 64 | NW_019014628.1 | 38 | 165 |
| 26 | NW_019014666.1 | 48 | 320 |
| 286 | NW_019014408.1 | 2 | 15 |

[a]Supercontigs mapping to the SDR

Appendix S5. Number of unique DEGs on LG1, the sex chromosome, and LG2-12, the autosomes, compared to the total number of genes in those regions.

| Genome location | DEGs | Total Genes | % of Total |
|---|---|---|---|
| LG1 (sex chromosome) | 217[a] | 2026 | 10.7% |
| LG2-12 (autosomes) | 1644 | 16880 | 9.7% |

[a]n.s., $P \geq 0.05$, Fisher's Exact Test

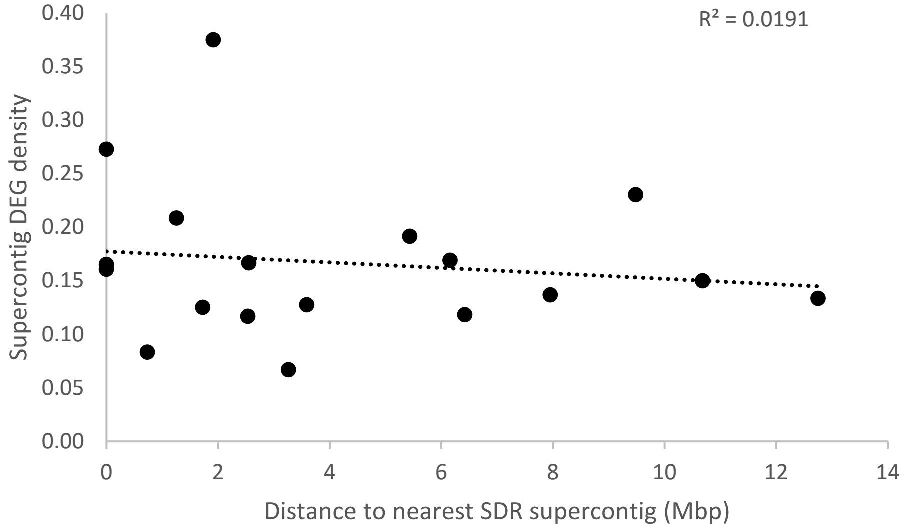


Appendix S6. Relationship between DEG density (number of DEGs in supercontig/total number of genes in that supercontig) to distance of that supercontig from the SDR. Supercontigs with fewer than 5 total genes were removed from the analysis (supercontigs 1066, 3603, 794, 2282, 438, 1076, 1124). There was no significant correlation between DE gene density and physical distance

Appendix S7. Location and expression bias of differentially expressed genes mapping to the X-PAR, X- and HSY-sequenced regions comparing male versus female early and late floral development.

| Gene ID | Linkage | Y Start | X Start | $Log_2$(Male/Female FPKM) | | Bias | Uniprot ID |
|---|---|---|---|---|---|---|---|
| | | | | Early | Late | | |
| 1 | X-PAR | - | 24110 | 3.25 | 3.72 | Male | CEPR2_ARATH Receptor protein-tyrosine kinase CEPR2 |
| 2 | X-PAR | - | 121657 | | **-0.70** | Female | TMN1_ARATH Transmembrane 9 superfamily 1 |
| 3 | X-PAR | - | 590054 | | **-1.44** | Female | CMBL_HUMAN Carboxymethylenebutenolidase homolog |
| 4 | X-PAR | - | 765046 | | 6.43 | Male | Y4598_ARATH Uncharacterized protein At4g06598 |
| 5 | X-PAR | - | 765509 | | **-1.04** | Female | Y4598_ARATH Uncharacterized protein At4g06598 |
| 6 | X-PAR | - | 775457 | | **-1.28** | Female | NDHT_ARATH NAD(P)H-quinone oxidoreductase subunit T, chloroplastic |
| 7 | X-PAR | - | 777444 | | **-3.50** | Female | ZHD4_ARATH Zinc-finger homeodomain protein 4 |
| 8 | X-PAR | - | 860049 | | **-2.92** | Female | FANCM_ARATH DEAD-box ATP-dependent RNA helicase FANCM |
| 9 | X-PAR | - | 897868 | | 1.82 | Male | CINV2_ARATH Alkaline/neutral invertase CINV2 |
| 10 | X-PAR | - | 992555 | | 0.53 | Male | GMCL1_MOUSE Germ cell-less protein-like 1 |
| 11 | X-PAR | - | 1025757 | | 1.05 | Male | SBT3H_ARATH Subtilisin-like protease SBT3.17 |
| 12 | X-PAR | - | 1300117 | | **-1.09** | Female | PIGQ_MOUSE Phosphatidylinitol N-acetylglucaminyltransferase subunit Q |
| 13 | X-PAR | - | 1823478 | 1.38 | 1.62 | Male | Y1343_ARATH G-type lectin S-receptor-like serine/threonine-protein kinase At1g34300 |
| 14 | X-PAR | - | 1827882 | | 4.92 | Male | PUB34_ARATH U-box domain-containing protein 34 |
| 15 | X | - | 2139631 | | **-1.46** | Female | POL3_DROME Retrovirus-related Pol polyprotein from transpon 17.6 |
| 16 | X | - | 2252636 | **-0.91** | | Female | EM506_ARATH Ankyrin repeat domain-containing protein EMB506, chloroplastic |
| 17 | X | - | 4114810 | | **-2.20** | Female | CYP38_ARATH Peptidyl-prolyl cis-trans isomerase CYP38, chloroplastic |
| 18 | X | - | 4808565 | | **-2.62** | Female | POL3_DROME Retrovirus-related Pol polyprotein from transospon 17.6 |
| 19 | X | - | 5111497 | | 2.28 | Male | ARF4_XENLA ADP-ribosylation factor 4 |
| 20 | Y | 870198 | - | 6.47 | Inf[a] | Male | MDAR4_ARATH Monodehydroascorbate reductase 4, peroxisomal |
| 21 | Y | 1110421 | - | Inf | Inf | Male | ARF1_ARATH ADP-ribosylation factor 1 |
| 22 | Y | 1118243 | - | Inf | Inf | Male | ARF1_ORYSJ ADP- ribosylation factor 1 |
| 23 | Y | 1206732 | - | 10.21 | Inf | Male | STKLD_ARATH Probable transcription factor At1g61730 |
| 24 | Y | 1876687 | - | Inf | Inf | Male | PGP1B_ARATH Phosphoglycolate phosphatase 1B, chloroplastic |
| 25 | Y | 3896873 | - | 4.06 | 3.76 | Male | IST1_RAT IST1 homolog |
| 26 | Y | 3903868 | - | 6.76 | 7.72 | Male | ATPD_ARATH ATP synthase subunit delta, chloroplastic |
| 27 | Y | 4003214 | - | Inf | Inf | Male | SVP_ARATH MADS-box protein SVP |
| 28 | Y | 4264884 | - | Inf | Inf | Male | CCD42_ARATH Cyclin-D4-2 |
| 29 | Y | 4291610 | - | 9.37 | Inf | Male | RK121_ARATH 50S ribosomal protein L12-1, chloroplastic |
| 30 | Y | 4449383 | - | Inf | Inf | Male | POLX_TOBAC Retrovirus-related Pol polyprotein from transposon TNT 1-94 |
| 31 | Y | 5123663 | - | 6.23 | 10.4 | Male | FRAY2_DICDI Serine/threonine-protein kinase fray2 |

[a]Inf = Infinity, result when there is no expression in females

Appendix S8. Location and expression bias of SDR-linked DEGs with X and Y gametologs comparing male versus female early and late floral development.

| X/Y Gene | | | Y Log$_2$(Male/Female FPKM) | | | X Log$_2$(Male/Female FPKM) | | | |
|---|---|---|---|---|---|---|---|---|---|
| Pair | Y Start | X Start | Early | Late | Bias | Early | Late | Bias | Uniprot ID |
| 1 | 241 | 4916151 | **-0.08** | **-1.43** | Female | **-1.31** | **-1.43** | Female | POLR2_ARATH Retrovirus-related Pol polyprotein from transposon RE2 |
| 2 | 1152056 | 2698684 | 11.84 | 11.03 | Male | | **-0.84** | Female | SSC14_ARATH Sister chromatid cohesion 1 protein 4 |
| 3 | 1293861 | 2843771 | 2.42 | 3.12 | Male | **-1.84** | | Female | BGL42_ARATH Beta-glucosidase 42 |
| 4 | 1381291 | 2930943 | 11.22 | 8.74 | Male | | **-0.89** | Female | HTSF1_HUMAN HIV Tat-specific factor 1 |
| 5 | 1542618 | 2967562 | Inf[a] | Inf | Male | | **-1.85** | Female | FAS1_ARATH Chromatin assembly factor 1 subunit FAS1 |
| 6 | 3794862 | 2048023 | 10.13 | 9.11 | Male | **-1.05** | **-1.34** | Female | SEC6_ARATH Exocyst complex component SEC6 |
| 7 | 3889362 | 1919675 | 5.65 | 6.03 | Male | | **-0.97** | Female | SERK1_ARATH Somatic embryogenesis receptor kinase 1 |
| 8 | 4375905 | 2660433 | 11.33 | Inf | Male | **-1.01** | **-0.71** | Female | LETM1_CHICK Mitochondrial proton/calcium exchanger protein |
| 9 | 4436403 | 2475393 | 9.62 | 7.00 | Male | | **-1.70** | Female | IP5PF_ARATH Type II inositol polyphosphate 5-phosphatase 15 |
| 10 | 4544077 | 4894559 | 4.12 | 3.81 | Male | **-1.48** | **-1.17** | Female | STY13_ARATH Serine/threonine-protein kinase STY13 |
| 11 | 4646038 | 4256601 | | Inf | Male | | 2.35 | Male | HAT5_ARATH Homeobox-leucine zipper protein HAT5 |
| 12 | 4651147 | 4229063 | 3.50 | 3.50 | Male | | **-1.47** | Female | MBD9_ARATH Methyl-CpG-binding domain-containing protein 9 |
| 13 | 4910346 | 4337804 | | 2.68 | Male | **-1.14** | | Female | DEGP1_ARATH Protease Do-like 1, chloroplastic |
| 14 | 5043844 | 4438681 | Inf | 4.11 | Male | | **-1.77** | Female | ALA8_ARATH Probable phospholipid-transporting ATPase 8 |
| 15 | 5137068 | 4568527 | 5.33 | Inf | Male | **-0.76** | | Female | PSMG3_DICDI Proteasome assembly chaperone 3 |
| 16 | 6971431 | 4624006 | INF | Inf | Male | **-1.62** | | Female | AGM1_ARATH Phosphoacetylglucosamine mutase |
| 17 | 7416822 | 3495662 | INF | Inf | Male | | **-1.45** | Female | ATM_ARATH Serine/threonine-protein kinase ATM |
| 18 | 7421623 | 3455492 | 6.43 | Inf | Male | **-1.23** | **-0.57** | Female | USPAL_ARATH Universal stress protein A-like protein |
| 19 | 7518078 | 3385187 | Inf | Inf | Male | **-1.14** | **-0.27** | Female | COG1_MOUSE Conserved oligomeric Golgi complex subunit 1 |
| 20 | 7638715 | 4948220 | | 1.59 | Male | | 1.64 | Male | FDH_SOLTU Formate dehydrogenase, mitochondrial |

[a]Inf = Infinity, result when there is no expression in females

Appendix S9. Location and expression bias of DEGs with X-PAR-, Y-PAR-, and Y-linked DEGs comparing female versus hermaphrodite late floral development.

| Gene ID | Linkage | Y Start | X Start | $Log_2$(Late Female/Late Herm FPKM) | Bias | Uniprot similarity |
|---|---|---|---|---|---|---|
| 1 | X-PAR | - | 142449 | **-2.88** | Herm | AAE5_ARATH Probable acyl-activating enzyme 5, peroxisomal |
| 2 | X-PAR | - | 203459 | **-3.23** | Herm | PMA10_ARATH ATPase 10, plasma membrane-type |
| 3 | X-PAR | - | 204697 | **-6.45** | Herm | BHLHW_PEA Basic helix-loop-helix protein A |
| 4 | X-PAR | - | 325791 | 3.96 | Female | PPR68_ARATH Pentatricopeptide repeat-containing protein At1g31920 |
| 5 | X-PAR | - | 392492 | **-7.24** | Herm | HA22J_ARATH HVA22-like protein j |
| 6 | X-PAR | - | 422884 | Inf[a] | Female | PTR1_ARATH Protein NRT1/ PTR FAMILY 8.1 |
| 7 | X-PAR | - | 480557 | **-2.52** | Herm | CIPKE_ARATH CBL-interacting serine/threonine-protein kinase 14 |
| 8 | X-PAR | - | 1570574 | **-3.45** | Herm | ZIP1_ARATH Zinc transporter 1 |
| 9 | Y-PAR | 21 | - | Inf | Female | YI31B_YEAST Transposon Ty3-I Gag-Pol polyprotein |
| 10 | Y-PAR | 605 | - | Inf | Female | NIN3_ORYSJ Neutral/alkaline invertase 3, chloroplastic |
| 11 | Y-PAR | 768 | - | 2.56 | Female | YI31B_YEAST Transpon Ty3-I Gag-Pol polyprotein |
| 12 | Y-PAR | 2578 | - | **-0.801** | Herm | ERDL6_ARATH Sugar transporter ERD6-like 6 |
| 13 | Y-PAR | 3464 | - | 2.60 | Female | FAFL_ARATH Protein FAF-like, chloroplastic |
| 14 | Y-PAR | 5552 | - | 0.595 | Female | PAPA3_CARPA Caricain |
| 15 | Y-PAR | 5959 | - | 3.70 | Female | TF26_SCHPO Transposon Tf2-6 polyprotein |
| 16 | Y-PAR | 7208 | - | **-7.57** | Herm | BXL1_ARATH Beta-D-xylidase 1 |
| 17 | Y-PAR | 11792 | - | **-4.05** | Herm | RIPK_ARATH Serine/threonine-protein kinase RIPK |
| 18 | Y-PAR | 12044 | - | 1.72 | Female | Y2921_ARATH Putative leucine-rich repeat receptor-like protein kinase At2g19210 |
| 19 | Y-PAR | 16332 | - | **-2.00** | Herm | GOT1_ARATH Vesicle transport protein GOT1 |
| 20 | Y-PAR | 18582 | - | **-3.05** | Herm | LSH3_ARATH Protein LIGHT-DEPENDENT SHORT HYPOCOTYLS 3 |
| 21 | Y-PAR | 30043 | - | **-3.55** | Herm | MARD1_ARATH Protein MARD1 |
| 22 | Y-PAR | 31408 | - | **-1.34** | Herm | PP443_ARATH Pentatricopeptide repeat-containing protein At5g62370 |
| 23 | Y-PAR | 125024 | - | **-4.85** | Herm | Y5248_ARATH G-type lectin S-receptor-like serine/threonine-protein kinase At5g24080 |
| 24 | Y-PAR | 163134 | - | Inf | Female | CHIA_ARATH Acidic endochitinase |
| 25 | Y | 960463 | - | **-6.83** | Herm | CYSEP_PHAVU Vignain |
| 26 | Y | 2086326 | - | **-1.27** | Herm | RMR1_ARATH Receptor homology region, transmembrane domain- and RING domain-containing protein 1 |
| 27 | Y | 2638401 | - | 1.63 | Female | CPS3B_ARATH Cleavage and polyadenylation specificity factor subunit 3-II |

[a]Inf = Infinity, result when there is no expression in hermaphrodites

Appendix S10. Expression analysis male X- and Y-gametologs and total male expression relative to total female expression.

| | | | Normalized expression[b] | | | | | |
|---|---|---|---|---|---|---|---|---|
| Stratum | ID[a] | Female Average FPKM | Female Total | Male X | Male Y | Male Total | Y:X | UNIPROT Similarity |
| 1 | 1 | 136.2 | 1 | 0.55 | 0.54 | 1.08 | 0.98 | SERK1_ARATH Somatic embryogenesis receptor kinase 1 |
| 1 | 2 | 97.3 | 1 | 0.89 | 0.99 | 1.88 | 1.10 | IST1_HUMAN IST1 homolog |
| 1 | 3 | 118.5 | 1 | 0.44 | 0.61 | 1.05 | 1.37 | SEC6_ARATH Exocyst complex component |
| 1 | 4 | 109.2 | 1 | 0.65 | 0.54 | 1.19 | 0.82 | |
| 1 | 5 | 343.5 | 1 | 0.36 | 0.15 | 0.51 | 0.42 | PGP1B_ARATH Phosphoglycolate phosphatase 1B, chloroplastic |
| 1 | 6 | 85.2 | 1 | 0.44 | 0.60 | 1.04 | 1.37 | IP5PF_ARATH Type II inositol polyphosphate 5-phosphatase 15 |
| 1 | 7 | 125.7 | 1 | 0.54 | 0.45 | 1.00 | 0.83 | LETM1_CHICK Mitochondrial proton/calcium exchanger protein |
| 1 | 8 | 104.5 | 1 | 0.63 | 0.64 | 1.26 | 1.02 | SSC14_ARATH Sister chromatid cohesion 1 protein 4 |
| 1 | 9 | 123.7 | 1 | 0.59 | 0.50 | 1.09 | 0.85 | HTSF1_HUMAN HIV Tat-specific factor 1 |
| 1 | 10 | 144.3 | 1 | 0.33 | 0.50 | 0.84 | 1.51 | FAS1_ARATH Chromatin assembly factor 1 subunit FAS1 |
| 1 | 11 | 140.6 | 1 | 0.68 | 0.64 | 1.32 | 0.94 | COG1_MOUSE Conserved oligomeric Golgi complex subunit 1 |
| 1 | 12 | 119.5 | 1 | 0.53 | 0.48 | 1.01 | 0.91 | USPAL_ARATH Universal stress protein A-like protein |
| 1 | 13 | 112.4 | 1 | 0.40 | 0.36 | 0.76 | 0.91 | ATM_ARATH Serine/threonine-protein kinase |
| 1 | 14 | 144.9 | 1 | 0.86 | 0.50 | 1.36 | 0.58 | ARF1_ORYSJ ADP-ribosylation factor 1 |
| 2 | 1 | 795.6 | 1 | 0.34 | 0.16 | 0.50 | 0.47 | TIO_ARATH Serine/threonine-protein kinase TIO |
| 2 | 2 | 267.7 | 1 | 0.97 | 0.29 | 1.26 | 0.30 | |
| 2 | 3 | 98.1 | 1 | 0.32 | 0.30 | 0.62 | 0.95 | CYP38_ARATH Peptidyl-prolyl cis-trans isomerase CYP38, chloroplastic |
| 2 | 4 | 26.9 | 1 | 1.52 | n.a[c] | n.a | n.a | SWT7B_ORYSJ Bidirectional sugar transporter SWEET7b |
| 2 | 5 | 72.0 | 1 | 0.59 | 0.60 | 1.19 | 1.03 | DEGP1_ARATH Protease Do-like 1, chloroplastic |
| 2 | 6 | 79.5 | 1 | 0.32 | 0.42 | 0.74 | 1.30 | CAHC_TOBAC Carbonic anhydrase, chloroplastic |
| 2 | 7 | 101.9 | 1 | 0.86 | 0.97 | 1.83 | 1.14 | |
| 2 | 8 | 99.2 | 1 | 0.39 | 0.59 | 0.98 | 1.53 | ALA8_ARATH Probable phospholipid-transporting ATPase 8 |
| 2 | 9 | 117.9 | 1 | 0.52 | 1.63 | 2.15 | 3.15 | |
| 2 | 10 | 106.0 | 1 | 0.69 | 0.52 | 1.21 | 0.76 | FRAY2_DICDI Serine/threonine-protein kinase fray2 |
| 2 | 11 | 118.6 | 1 | 0.76 | 0.61 | 1.37 | 0.80 | PSMG3_DICDI Proteasome assembly chaperone 3 |
| 2 | 12 | 112.9 | 1 | 0.45 | 0.88 | 1.32 | 1.97 | AGM1_ARATH Phosphoacetylglucosamine mutase |
| 2 | 13 | 70.6 | 1 | 0.32 | n.a | n.a | n.a | AS2_ARATH Protein ASYMMETRIC LEAVES 2 |
| 2 | 14 | 62.8 | 1 | 0.41 | 0.89 | 1.30 | 2.15 | |
| 2 | 15 | 309.3 | 1 | 0.28 | 0.11 | 0.40 | 0.39 | |
| 2 | 16 | 180.7 | 1 | 0.41 | 0.22 | 0.63 | 0.53 | STY13_ARATH Serine/threonine-protein kinase STY13 |

[a]Gene ID corresponds to ID in Fig. 4; [b]Expression normalized relative to total Female expression; [c]n.a. not applicable due to no FPKM reading.

Appendix S10, cont.

| | | | Normalized expression[b] | | | | | |
|---|---|---|---|---|---|---|---|---|
| Stratum | ID[a] | Female Average FPKM | Female Total | Male X | Male Y | Male Total | Y:X | UNIPROT Similarity |
| 3 | 1 | 143.2 | 1 | 0.24 | 0.39 | 0.63 | 1.66 | SAMH1_DICDI Deoxynucleoside triphosphate triphosphohydrolase |
| 3 | 2 | 44.9 | 1 | 2.04 | 1.19 | 3.23 | 0.58 | NADB_ARATH L-aspartate oxidase, chloroplastic |
| 3 | 3 | 27.9 | 1 | 2.65 | 2.75 | 5.40 | 1.04 | ARF_DUGJA ADP-ribosylation factor |
| 3 | 4 | 190.6 | 1 | 0.67 | 0.45 | 1.12 | 0.66 | |
| 3 | 5 | 45.9 | 1 | 0.75 | 1.30 | 2.05 | 1.73 | RSH3C_ARATH Probable GTP diphosphokinase RSH3, chloroplastic |
| 3 | 6 | 59.9 | 1 | 0.57 | 1.50 | 2.06 | 2.65 | XPO1A_ARATH Protein EXPORTIN 1A |
| 3 | 7 | 47.0 | 1 | 0.65 | 2.18 | 2.83 | 3.38 | NDUS6_ARATH NADH dehydrogenase [ubiquinone] iron-sulfur protein 6 |
| 3 | 8 | 62.3 | 1 | 0.64 | 1.38 | 2.02 | 2.16 | PP399_ARATH Pentatricopeptide repeat-containing protein At5g27270 |
| 3 | 9 | 31.7 | 1 | 1.44 | n.a[c] | n.a | n.a | |
| 3 | 10 | 49.3 | 1 | 1.16 | 1.19 | 2.35 | 1.03 | XYLL2_ARATH D-xylose-proton symporter-like 2 |

[a]Gene ID corresponds to ID in Fig. 4; [b]Expression normalized relative to total Female expression; [c]n.a. not applicable due to due to no FPKM reading
